# GeneSurfer Enables Transcriptome-wide Exploration and Functional Annotation of Gene Co-expression Modules in 3D Spatial Transcriptomics Data

**DOI:** 10.1101/2024.07.05.602230

**Authors:** Chang Li, Julian Thijssen, Thomas Kroes, Ximaine van der Burg, Louise van der Weerd, Thomas Höllt, Boudewijn Lelieveldt

## Abstract

Gene co-expression provides crucial insights into biological functions, however, there is a lack of exploratory analysis tools for localized gene co-expression in large-scale datasets. We present GeneSurfer, an interactive interface designed to explore localized transcriptome-wide gene co-expression patterns in the 3D spatial domain. Key features of GeneSurfer include transcriptome-wide gene filtering and gene clustering based on spatial local co-expression within transcriptomically similar cells, multi-slice 3D rendering of average expression of gene clusters, and on-the-fly Gene Ontology term annotation of co-expressed gene sets. Additionally, GeneSurfer offers multiple linked views for investigating individual genes or gene co-expression in the spatial domain at each exploration stage. Demonstrating its utility with both spatial transcriptomics and single-cell RNA sequencing data from the Allen Brain Cell Atlas, GeneSurfer effectively identifies and annotates localized transcriptome-wide co-expression, providing biological insights and facilitating hypothesis generation and validation.

## Introduction

The rapid emergence of single-cell RNA sequencing (scRNA-seq) and Spatial Transcriptomics (ST) has led to routine generation of high-dimensional, spatially resolved datasets at the whole-brain scale^1-3^. These datasets offer high informational content but require scalable computational methods for effective data exploration.

From spatially sampled bulk sequencing we already know that spatially co-varying genes tend to be functionally related^4^. The “guilt by association” principle suggests that genes which are co-expressed are likely to share function roles, especially within transcriptomically similar cells.

Methods for grouping genes into co-expression modules include clustering, matrix decomposition, network inference-based approaches and others^5^. The most common approach is to employ clustering algorithms for gene set identification. However, one major limitation is that clustering methods group genes using the entire dataset as input. It might overlook spatially localized co-expression patterns within specific cell subsets, since genes can be locally co-expressed at certain locations in the brain but not at others. Thus, methods that can detect spatially localized co-expression could provide biological specificity of co-expression patterns. Among the various module detection methods, biclustering is able to go beyond this limitation by simultaneously capturing gene co-expression and sample subsets across the data matrix^6,7^. But similar to other module detection methods, biclustering algorithms are script-based and require predefined parameters^5^. This introduces uncertainty and leads to the need for testing across different settings. Therefore, there still exists a gap to identify the local gene co-expression in a flexible and interactive manner for the user.

Besides methods that directly identify local gene co-expression, we can also apply the clustering algorithm to a subset of a specific cell type, to compute co-expression within transcriptomically similar cells. However, cell clustering algorithms divide cells into disjoint classes according to transcriptional similarity, binarizing potentially gradual cell type transitions and gradients into distinct subtypes^3^. This process requires the user to decide beforehand which level of cluster resolution is of interest, limiting the ability to explore the complete parameter space. Currently, there is a lack of approaches that allow for the identification of local spatial co-expression at user-defined cell-type resolution, offering a continuum approach to cell transcriptome similarities.

ST techniques measure gene expression information of tens to hundreds of genes in individual cells, while preserving their spatial context in the tissue^2^. This enables the visualization of gene expression in the spatial domain. Despite significant improvements in the transcriptome resolution in recent years^2^, a primary limitation remains that spatially resolved transcriptomics captures far fewer genes compared to scRNA-seq. In contrast, scRNA-seq aims to capture a detailed transcriptomic profile, but does not provide spatial context. Integrative tools have been developed to facilitate the visualization of gene expression in ST data or to represent scRNA-seq data in reduced dimensions^8-10^, such as through t-distributed stochastic neighbor embedding (t-SNE)^11^ and uniform manifold approximation and projection (UMAP)^12^. Additionally, computational imputation methods and machine learning models are actively being researched to integrate scRNA-seq data to expand the gene dimensionality of ST data^13-16^. However, most of these methods are static, and interactive visualization softwares for visualizing large-scale scRNA-seq data in the 3D spatial domain are lacking.

Subsequent steps of co-expression identification usually include visualization and functional interpretation. Gene set enrichment analysis allows us to derive molecular insights with interpretable terms, such as known biological pathways and functions^17^. Enrichment frameworks such as ToppGene^18^ and gProfiler^19^ offer Application Programming Interface (API) access to integrate enrichment analysis into analytical tools.

In this work, we introduce GeneSurfer: an interactive interface for exploring localized transcriptome-wide gene co-expression patterns in the 3D spatial domain. The major contributions of our work are: 1) transcriptome-wide gene filtering and gene clustering based on spatial co-expression within transcriptomically similar cell populations, 2) real-time, interactive cell selection and multi-slice 3D rendering of average expression of gene clusters, and 3) on-the-fly Gene Ontology (GO) term annotation of co-expressed gene sets. In addition, we offer multiple linked views for interactive visualization of individual genes or gene co-expression in the spatial domain at each stage of the exploration. We demonstrate the utility of GeneSurfer using both ST and scRNA-seq data from the Allen Brain Cell (ABC) Atlas^1^.

## Results

### GeneSurfer Overview

By offering multiple linked views of cells and genes, we aim to provide the user with an interactive interface to explore local gene expression patterns in the spatial domain, and to gain potential biological insights from the identified spatial patterns (Supplemental Movie 1). We support the input of ST datasets, and additionally we also project the averages of transcriptome-wide gene expression obtained in scRNA-seq data onto the spatial domain by matching annotations (Figure 1B). This requires that both the ST and the scRNA-seq are classified in the exact same cell type taxonomy.

**Figure 1.**
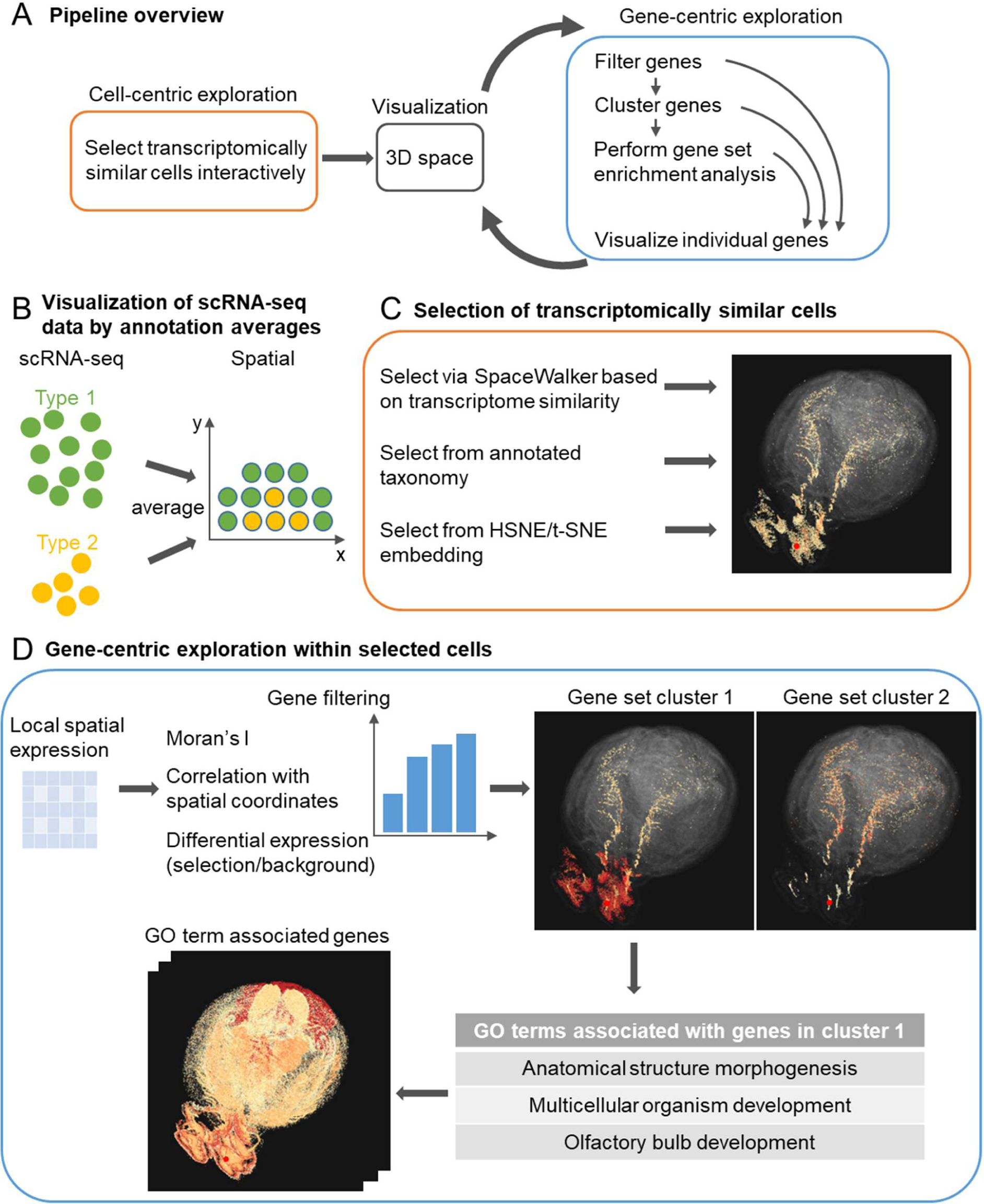
GeneSurfer is an interactive interface for exploring local gene expression patterns in the spatial domain. **A.** Overview of the proposed workflow. Users select transcriptomically similar cells and visualize them in 3D. Genes are filtered and then clustered based on their expression within the selected cells. GO terms are retrieved by selecting a gene set cluster, and individual genes can be visualized in 2D and 3D. **B.** scRNA-seq expression averages of the annotations are projected in the spatial domain, allowing visual exploration of transcriptome-wide gene expression in 3D space. **C.** Users can interactively select transcriptomically similar cells using SpaceWalker, annotations from the data taxonomy, or through HSNE or t-SNE embeddings. **D.** After selecting a subset of cells, genes are filtered based on their local expression within the selected cells. Three metrics are available for gene filtering: Moran’s I, Pearson correlation between gene expression and corresponding spatial coordinates and differential expression between the selection and the whole data. Filtered genes are displayed in a bar chart, and are clustered into groups. Associated GO terms of the selected gene cluster are retrieved and displayed in a table. Views in GeneSurfer are linked to enable an interactive manner of exploration.

An overview of the proposed methodology is shown in Figure 1A. Our approach begins with a cell-centric exploration, where users select cells of interest that have similar gene expression profiles through one of the three selection methods (Figure 1C). Firstly, users can select a cell type annotation from the annotated data taxonomy. Secondly, users can use our previous work SpaceWalker^20^ to interactively identify cells with transcriptomically similar profiles at user-defined cell locations. And thirdly, users can re-embed the data on-the-fly using Hierarchical Stochastic Neighbor Embedding (HSNE)^21^ or t-SNE^11^ and select cell clusters from the embedding. As such, this initial step focuses on selecting cell populations of interest for further analysis.

Once similar cells are selected, this workflow instantaneously transitions to gene-centric exploration (Figure 1D), allowing for on-the-fly analysis of genes within the selected cells. Prior to gene set clustering, gene filtering is performed based on user-defined spatial metrics within the cell selection (Moran’s I statistics^22^ or the correlation of gene expression and coordinate axes in a standardized Common Coordinate Frame (CCF)). In addition, a non-spatial gene filter can be applied to identify genes highly expressed in the selected cells.

Based on the expression profiles of the selected cells, genes are grouped into clusters with similar local spatial patterns using hierarchical clustering^23^. The average expression of genes in each cluster is shown in subviews to provide an overview of the expression patterns of the gene clusters. Gene set enrichment analysis is performed instantaneously on the user-selected gene cluster to provide insights into the potential biological functions or pathways associated with the genes. The output of enrichment analysis is presented in a table listing the statistically significant GO term annotations, their corresponding p-values, and associated gene symbols.

Overall, GeneSurfer allows a closed-loop, interactive exploration of gene expression in a user-defined subset of cells.

### Projection of scRNA-seq annotation averages enables visual exploration of transcriptome-wide gene expression in 3D space

To evaluate the projection of annotation averages from scRNA-seq data into the spatial domain based on label correspondence, we used the ABC Atlas^1^. The ABC Atlas is a mouse whole-brain transcriptomic cell type atlas, consisting of an annotated scRNA-seq dataset with ∼4 million cells and more than 30k genes and an annotated 3D ST dataset with ∼3.9 million cells and 500 genes using multiplexed error-robust fluorescence in situ hybridization (MERFISH). The fine-grained cell-type annotations (> 5000 cell types) in the ABC Atlas make it an appropriate benchmark dataset for validating our tool, while demonstrating the scalability and discovery potential of the proposed approach. The annotation level “Cluster” is used for the projection.

To validate the 3D spatial projection of scRNA-seq annotation averages, we performed two quantitative experiments, comparing the spatial gene expression patterns of the 500 genes that are measured in both the MERFISH and scRNA-seq data. First, to establish the validity of replacing expression in the 3D spatial data with expression averages of the corresponding annotation from scRNA-seq data, we calculated the Pearson correlation at the cluster annotation level between the scRNA-seq annotation averages and the measured MERFISH data for each gene available in the MERFISH data. Figure 2A shows that the mean expression levels of genes in the clusters are highly correlated between the scRNA-seq clusters and the MERFISH clusters. Second, we calculated the spatial Pearson correlation between the 3D projection of the scRNA-seq annotation averages and the actual MERFISH measurement at the cell level in the 3D spatial domain. Figure 2B shows that the spatial correlations at the cell level are lower than those at the cell cluster level. Thus we further inspected the individual gene expression by visually comparing the spatial patterns of scRNA-seq annotation averages and the MERFISH measurement. Several genes exhibit high correlation, particularly in genes with widespread and distinctive expression in the whole brain (e.g. *Cbln1* in Figure 2C). The relatively low correlations might be due to the fact that the correlation at the cell level is calculated for the whole brain (∼4 million cells). A gene that expresses in the whole brain in a scattered way, or a gene that only expresses highly in a small spatially localized hotspot would result in a low correlation coefficient, for example *Lsp1* and *Foxa2* (Figure 2C). However, the main structure of the gene expression is preserved by label correspondence of scRNA-seq annotation averages, despite some instances of low correlation values.

**Figure 2.**
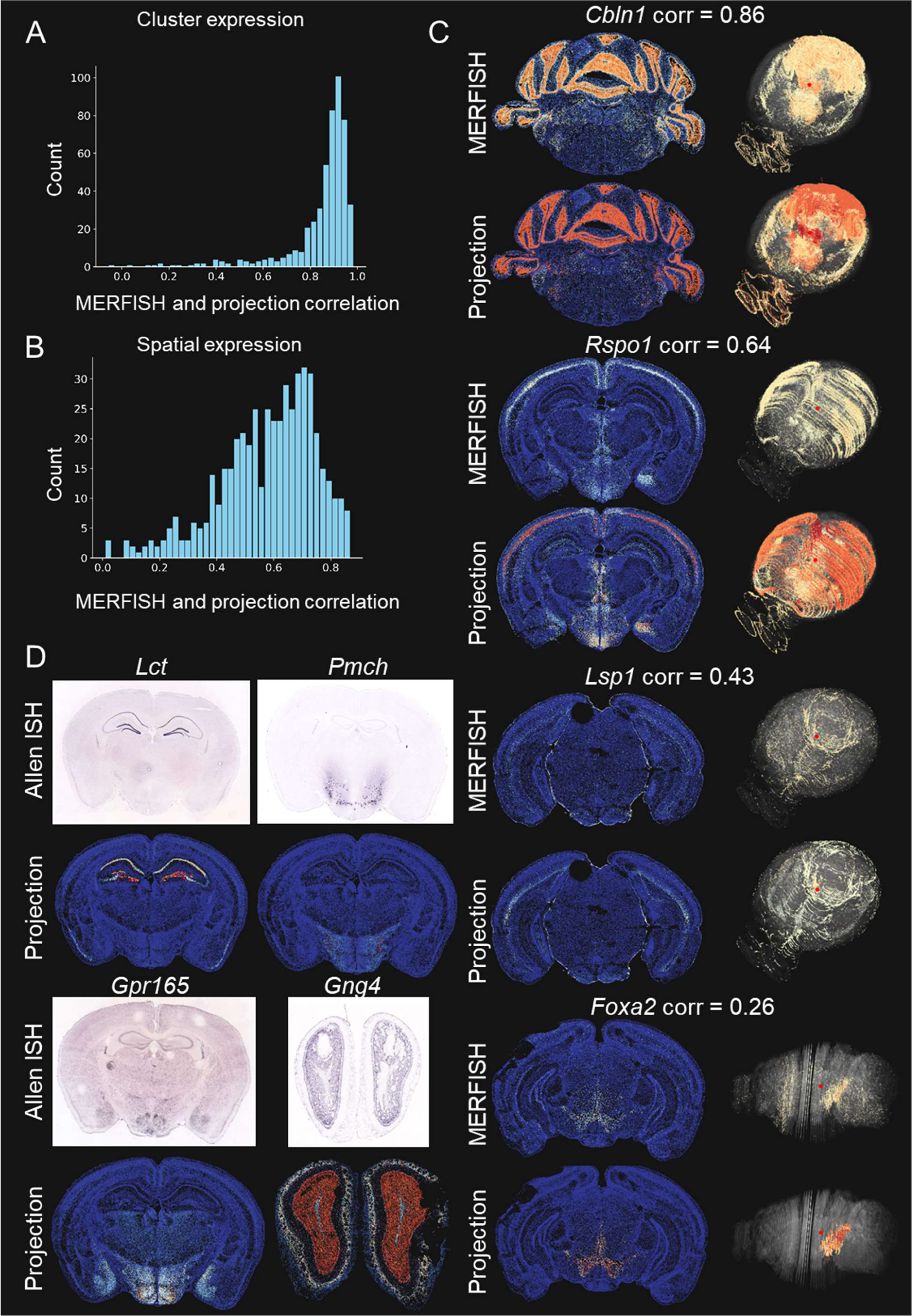
Comparison of projection of scRNA-seq annotation averages with the MERFISH measurement and Allen in situ hybridization data. **A.** The distributions of Pearson correlation coefficients between MERFISH measurement and scRNA-seq annotation averages in cell clusters. **B.** The distributions of spatial Pearson correlation coefficients between MERFISH measurement and projection of scRNA-seq annotation averages across all cells. **C.** Examples of spatial gene expression patterns and their spatial correlations of MERFISH measurement and projection of scRNA-seq annotation averages. **D.** Examples of spatial gene expression patterns from Allen Brain Atlas in situ hybridization data and projection of scRNA-seq annotation averages. The Allen Brain Atlas in situ hybridization data are taken from https://mouse.brain-map.org/^4^.

To evaluate the projection of scRNA-seq annotation averages for the genes not measured in the MERFISH, we compared the spatial gene expression between projection of the scRNA-seq annotation averages and the Allen Brain Atlas in situ hybridization data^4^. Projection of the scRNA-seq annotation averages show excellent agreement with the Allen Brain Atlas in situ hybridization data (Figure 2D).

### GeneSurfer identifies spatial co-expression within user-defined cell populations

GeneSurfer aims to identify genes with distinct spatial co-expression within interactively selected transcriptomically similar cell populations. To validate the gene filtering based on spatial expression, and the subsequent gene clustering, we next investigated whether GeneSurfer can identify distinct local spatial co-expression using different gene filtering options.

GeneSurfer provides three gene filtering options to enhance the further gene-centric exploration within selected cells: 1) correlation with a spatial coordinate axis in the CCF to identify genes exhibiting a spatial gradient along the anatomical brain axes, 2) Moran’s I^22^ to detect genes with non-random spatial expression patterns within the cell selection, and 3) differential expression between foreground and background cells to identify genes highly expressed in the selection. Filtered genes are clustered into groups, revealing potentially distinct spatial co-expression across the brain anatomy. Details on these gene filtering metrics and gene clustering are described in the Methods section.

Figure 3 provides visual examples of the gene filtering and clustering results obtained with GeneSurfer using the projection of scRNA-seq annotation averages. The spatial correlation filters on the anterior-posterior or dorsal-ventral axis identify genes with spatial gradients along an anatomical axis in the CCF. We first selected the IT-ET Glut class from the data taxonomy to explore the spatial patterns within the excitatory neurons in the cortex. Genes with local gradients on the dorsal-ventral axis were filtered out and were clustered into two groups with distinct gradients: dorsal-ventral gradient and ventral-dorsal gradient (Figure 3A). Then we drilled into L5 IT, L6 IT and L5/6 IT Glut subclasses to explore the potential anterior-posterior gradients at a more refined cell-type level: excitatory neurons in the deeper cortical layers. Genes that exhibit an anterior-posterior gradient or posterior-anterior gradient within L5/L6 IT neurons were identified and subsequently clustered into two gene clusters (Figure 3B). The Moran’s I filter highlights genes with patterns that significantly deviate from randomness, suggesting potential functions within the spatial context. Figure 3C shows the two different expression patterns identified within the OB-IMN GABA class, where genes are filtered by the Moran’s I statistic. Genes in cluster 1 show a high expression in the subgranular zone, subventricular zone and rostral migratory stream (SGZ, SVZ and RMS), which might be related to neuron differentiation and cell proliferation^1^, including marker genes such as *Prox1*. Genes in cluster 2 are highly expressed in the olfactory bulb, with decreasing expression toward the posterior aspect of the brain, which might be related to mature olfactory bulb neurons.

**Figure 3.**
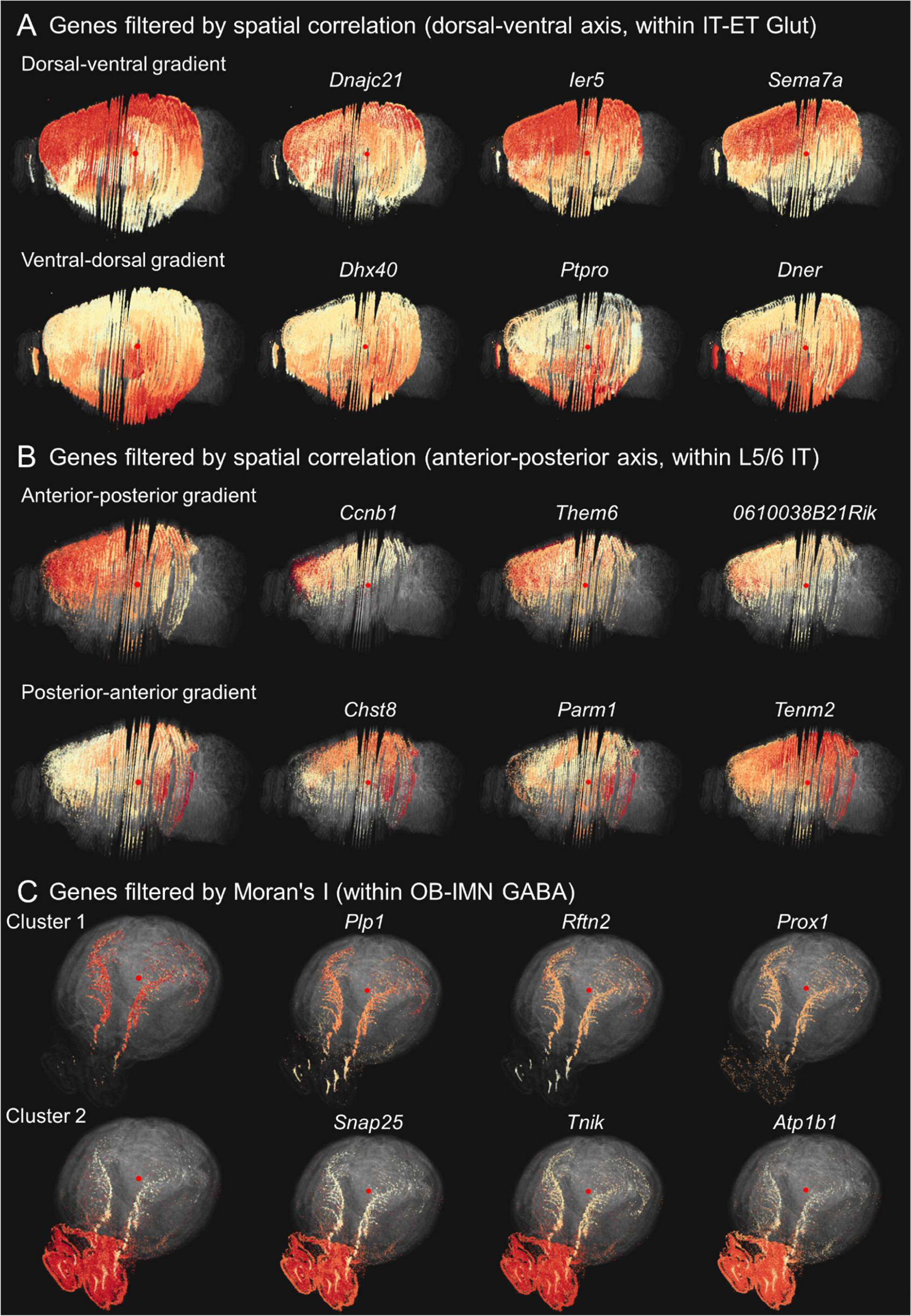
Example results of gene filtering using different metrics and subsequent gene clustering. **A.** Genes filtered by spatial correlation on the dorsal-ventral axis within IT-ET Glut neurons. Filtered genes were clustered into two groups: genes with positive spatial correlation show a superior-inferior gradient, with the reverse for genes with negative spatial correlation. **B.** Genes filtered by spatial correlation on the anterior-posterior axis within L5 IT, L6 IT and L5/6 IT neurons. Filtered genes were clustered into two groups: genes with a high positive correlation display an anterior-posterior gradient, with the reverse for genes with negative spatial correlation. **C.** Genes filtered using Moran’s I within OB-IMN GABA class exhibit non-random expression patterns and are clustered into two groups. The first column in panels A, B, and C displays average expression of genes in each cluster. The gene expression patterns shown are within the selected subset and are color-coded based on normalized values within the subset. (Parameters: 100 genes, 2 clusters)

These examples demonstrate GeneSurfer’s utility in filtering genes with spatial expression and clustering genes to identify local co-expression, and also guide users in selecting the most appropriate gene filtering metrics based on their specific research questions.

The differential expression filter is used to identify genes that are predominantly expressed in the selection. This is designed to explore cell-type specific gene expression which can subsequently be inspected in the spatial domain, which is demonstrated in the next section.

### Interactive exploration with GeneSurfer confirms known biological function

In this section, we demonstrate the interactive exploration workflow with GeneSurfer on the whole transcriptome by validating its findings against known biology.

To evaluate the exploration workflow of GeneSurfer together with SpaceWalker, we demonstrate the workflow by exploring cells on a 2D brain slice. GO term annotations were then retrieved for these cell populations and compared to known biological pathways for these cells. When a seed cell location was selected in SpaceWalker, transcriptomically similar cells were selected by flood-fill^20^ on the fly. Subsequently in GeneSurfer, 50 genes were filtered out employing the differential expression filter that compares gene expression between selection and background cells. These genes were then clustered into two groups based on their local expression. GO term annotations were then instantaneously retrieved for gene function annotations. Figure 4 presents the results of selecting cells at four spatial locations that are close to each other spatially but transcriptomically and functionally distinct, annotated as subclass astrocytes (Astro-TE NN), oligodendrocytes (Oligo NN), pericytes (Peri NN) and microglia (Microglia NN). Genes associated with corresponding functions were identified and associated GO terms were retrieved, demonstrating that SpaceWalker identified transcriptomically similar cells and GeneSurfer discovered differentially expressed genes within this selected subset of cells, reflecting the biological function of these cells. These findings confirm the capability of GeneSurfer with SpaceWalker to analyze cellular components and gene expression in an explorative manner.

**Figure 4.**
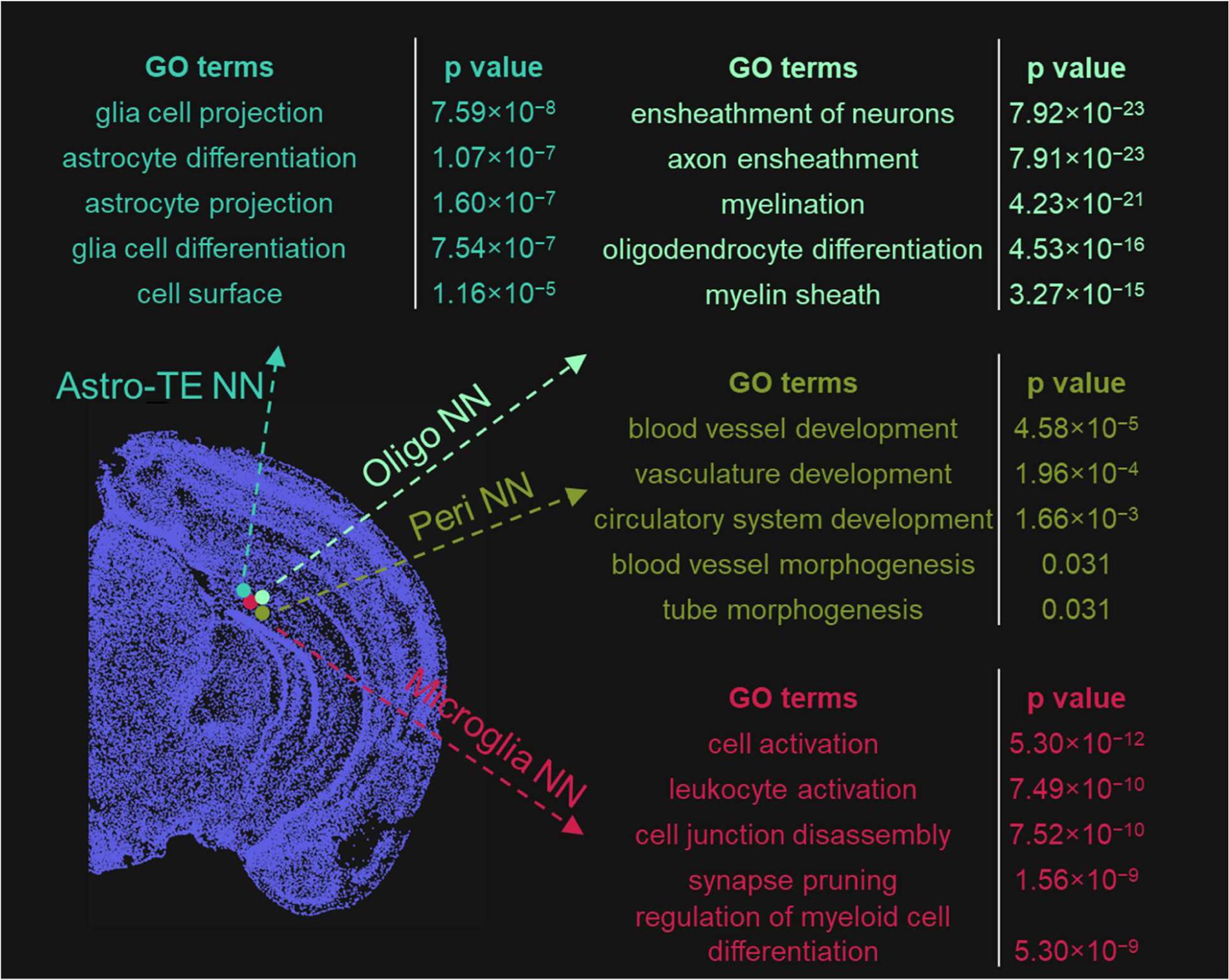
Transcriptome-wide exploration using SpaceWalker and GeneSurfer retrieved cell-type specific GO term annotations. Hovering over the 2D mouse brain section allows for selecting different cells at close-by locations (annotated as Astro-TE NN, Oligo NN, Peri NN and Microglia NN). Transcriptomically similar cells are identified by SpaceWalker and genes are filtered by the differential expression filter within the transcriptomically similar cells. For each selected cell location, GO terms associated with the filtered genes are retrieved, with the figure presenting the five GO terms having the lowest p-values. scRNA-seq annotation averages serve as input for gene filtering and clustering. GO terms for scRNA-seq genes are retrieved by ToppGene without a background gene set. All p-values in this paper have been adjusted using the Bonferroni correction. (GeneSurfer: differential expression filter, 50 genes, 2 clusters; SpaceWalker: flood nodes = 10, flood steps =10)

It is important to note that cell annotations were only used to validate the results with the known annotation, but the methodology does not use these prior annotations in the gene-centric process and annotations are not required for this workflow. Annotations were only used to project scRNA-seq data to the spatial domain, not in the gene-centric exploration with selecting cells with SpaceWalker. Thus, ST data without annotations can also be explored using this workflow.

In addition to selecting transcriptomically similar cells through SpaceWalker and the annotation taxonomy, as GeneSurfer is built on top of the ManiVault API^24^, GeneSurfer seamlessly integrates with other ManiVault plugins to extend functionality, such as on-the-fly dimensionality reduction. Users can re-embed the whole dataset or a subset of the data, color the embedding using existing annotations, and select cells directly from the embedding to further explain the dataset in the gene-centric workflow.

Although this Genesurfer workflow is extremely powerful, the gene-centric clustering may be affected by the artifacts in the preprocessing of the MERFISH data. We show here an example of endothelial cells, which are annotated as three distinct clusters in the ABC Atlas^1^. The re-embedded HSNE map using the MERFISH data reveals additional substructures within the Endo NN_1 cluster (Figure 5A). Selecting points of interest in the HSNE embedding, we observed spatially distinct structures within the Endo NN_1 cluster (Figure 5B). To further investigate whether this suggests additional cellular heterogeneity within the Endo NN_1 cluster in the MERFISH data, we performed differential expression analysis between these substructures and the Endo NN_1 cluster. We then analyzed the scRNA-seq data to inspect cell-specificity of the top differentially expressed genes. We found that each substructure was characterized by a small set of differentially expressed genes that were in most cases expressed in other cell types surrounding endothelial cells, but with region-specific abundance. Based on these observations, we hypothesize that the observed substructures in the spatial data could be caused by a spillover effect of gene transcripts between cell types during the segmentation of (small) endothelial cells. Thus, GeneSurfer enables users to examine the annotated data, and interactively differentiate potential cellular heterogeneity from underlying artifacts inherently present in the data.

**Figure 5.**
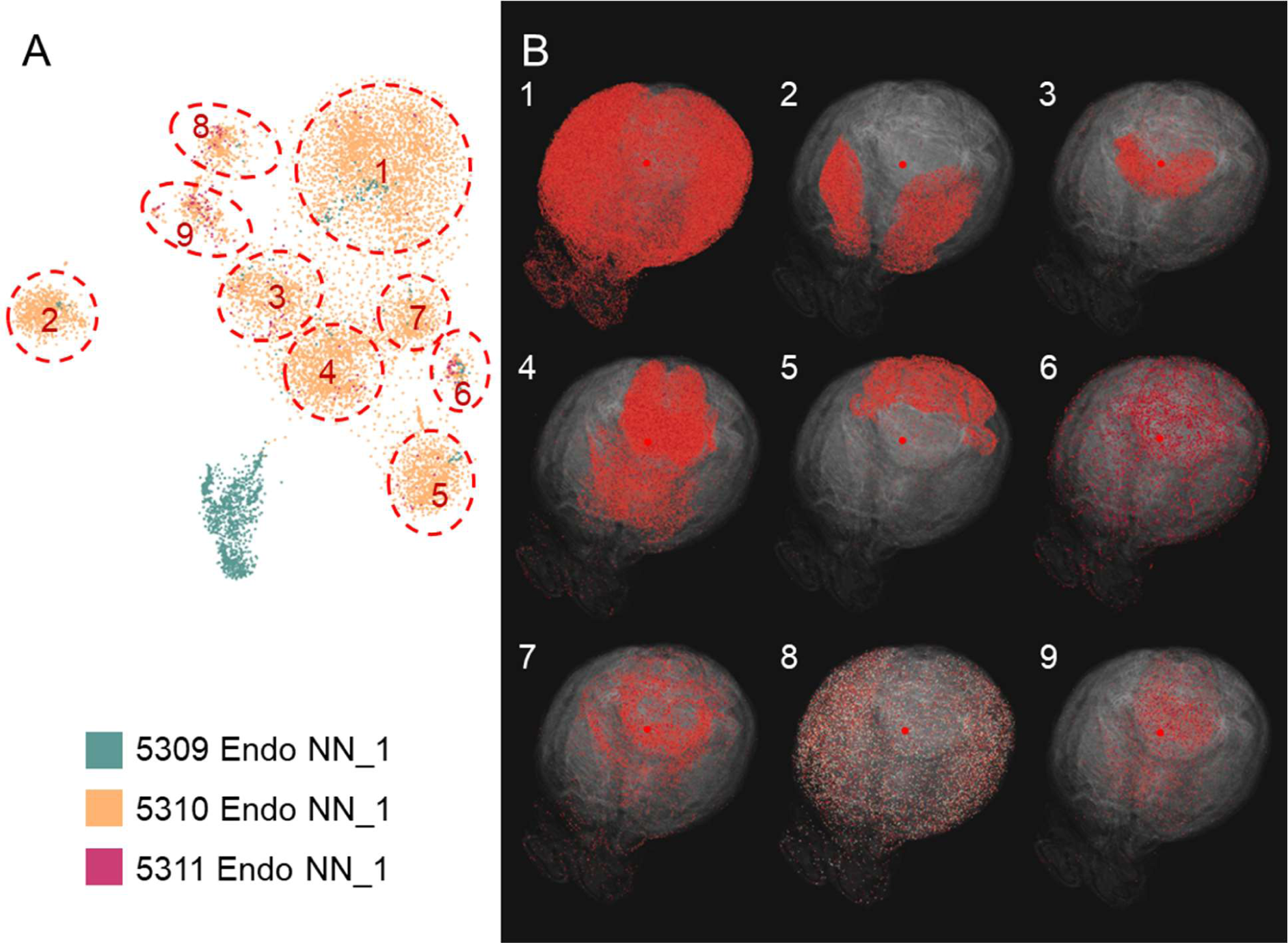
Demonstration of exploring endothelial cells with HSNE re-embedding. **A.** HSNE embedding of endothelial cells from the MERFISH data, colored by cluster annotation^1^. Substructures in the embedding map are circled and numbered on the map. B. The corresponding substructures marked in (A) are visualized in the 3D spatial domain, displaying distinct spatial structures. Inspection of the top differentially expressed genes and comparison with the scRNASeq data lead to the conclusion that these sub-clusters are most likely an artifact, due to spillover from surrounding cells.

## Discussion

Recent advances in transcriptomics have led to the need for computational tools to explore large-scale datasets. Computational approaches have been developed to identify co-expressed genes, indicating their functional roles. However, current methods fall short in providing an exploratory tool that can interactively identify spatial co-expression, particularly in a local cellular subspace.

Here, we present GeneSurfer, an interactive interface to explore localized transcriptome-wide gene co-expression in the 3D spatial domain. It allows users to dynamically explore local co-expressed genes within the user-selected cells and on-the-fly GO term annotation of these co-expressed gene clusters. We also support an approximate visualization of scRNA-seq imputed gene expression onto the spatial domain, enabling the user to interactively explore both ST and scRNA-seq gene expression in the 3D spatial domain. In addition, we offer multiple linked views for interactively investigating individual genes or gene co-expression in the spatial domain at each stage of the exploration. By integrating multiple ways to select cells of interest, three gene filtering options and various user-defined clustering and visualization parameters, GeneSurfer enables user-tailored exploration paths.

GeneSurfer has novel capabilities in comparison to existing approaches. Compared with existing algorithms for identifying co-expression modules in the whole data^5^, we specifically target localized spatial co-expression within transcriptomically similar cells in an interactive manner. Secondly, unlike computational methods to integrate scRNA-seq with ST data^13-16^, which primarily focus on data integration, our tool extends the functionality to provide transcriptome-wide gene analysis and visualization in the 3D spatial domain by matching annotations. Moreover, while some integrative toolboxes offer interactive visualization^8-10^, they do not support interactive and localized gene-centric on-the-fly computational analysis as our tool does. While our previous work SpaceWalker^20^ focuses on cell-centric exploration and expression gradient detection, GeneSurfer focuses on a gene-centric analysis approach to explore transcriptome-wide gene co-expression within selections of similar cells. Upon selecting cells, GeneSurfer performs the analysis on-the-fly, providing a responsive and interactive experience on a mid-range PC workstation.

We validated the functionality of GeneSurfer using the 3D whole-brain mouse dataset ABC Atlas. We demonstrated that GeneSurfer is able to identify and annotate local gene co-expression within selected cells, providing biological insights in real-time during exploration. We used the projection of annotation averages from scRNA-seq data onto the spatial domain to enable transcriptome-wide exploration, demonstrating its validity when both scRNA-seq and ST datasets are annotated and available. Additionally, GeneSurfer can work with unannotated ST datasets to help with hypothesis generation, and with the design of targeted validation experiments. It is compatible with both 3D and 2D datasets.

Overall, GeneSurfer offers an interactive visual interface for exploring local gene co-expression and their functional annotation.

## Acknowledgements

This work received financial support from NWO Gravitation project BRAINSCAPES: A Roadmap from Neurogenetics to Neurobiology (NWO: 024.004.012); the NIH Brain Initiative Cell Atlas Network BICAN (HMBA: UM1MH130981); NWO TTW project 3DOMICS (NWO: 17126) and Alzheimer Netherlands project WE.03-2022-15.

## Author contributions

C.L., T.H. and B.L. conceptualized the project. C.L., J.T. and T.K. implemented the software. C.L., X.v.d.B., L.v.d.W. and B.L. performed the analyses and interpreted the data. C.L. and B.L. wrote the manuscript with assistance and approval from all authors.

## METHODS

## RESOURCE AVAILABILITY

### Materials availability

This study did not generate new unique reagents.

### Data and code availability

- This paper analyzes existing, publicly available data.
- GeneSurfer is implemented in C++ as a plugin of the ManiVault plugin system^24^ for visual analytics application building. All original code, along with the Windows installer and a system state file containing the data, plugins, and GUI configurations, is publicly available at https://github.com/ManiVaultStudio/GeneSurfer. Installers for MacOS and Linux will be made available at https://www.manivault.studio in the future.

### Method details

#### Data

ABC Atlas data was downloaded from https://allen-brain-cell-atlas.s3.us-west-2.amazonaws.com/index.html. We used both the MERFISH data and scRNA-seq data for the analysis in this paper.

#### Projecting scRNA data to ST domain

To visualize scRNA-seq data in the spatial domain, we utilized the matching cell annotations in the ST and scRNA-seq data. After loading both ST and scRNA-seq datasets, the user selects an annotation dataset to guide the alignment of the two datasets. The system checks whether both the ST and scRNA-seq datasets contain the required metadata for the selected annotation. The average expression level is then calculated for each gene of each annotation category in the scRNA-seq data. These calculated average expressions are assigned to the spatial coordinates in the ST data based on their corresponding annotations, replacing the actual measured MERFISH expressions at that location. In this way, we get an approximate gene expression matrix from scRNA-seq data with spatial coordinates, allowing for visualizing and analyzing the projected scRNA-seq gene sets in 2D and 3D spatial domain. In this manuscript, the annotation level “Cluster” is used for the ABC Atlas.

The prior condition is that an annotated ST dataset and an annotated scRNA-seq dataset of the same region are required, where the latter covers major annotation types in the ST dataset. We assume that the annotations in the datasets are fine-grained enough that the average expression of each annotation class is a representative approximation for that annotation class.

Since some of the scRNA-seq datasets can be prohibitively large, ingesting all of the data in memory to compute annotation averages may result in memory overflows or real-time computational difficulties. Users have the option to load a pre-computed average expression matrix instead of computing the matrix on-the-fly; this obviates the need to ingest the entire scRNAseq data to compute cluster statistics.

In summary, we bridge the gap in gene coverage between scRNA-seq and ST data by directly projecting average expression based on fine-grained annotations, to provide an approximate visualization of scRNA-seq data in the spatial domain.

#### Selection of transcriptomically similar cells

GeneSurfer focuses on the gene-centric exploration within a subset of cells with similar transcriptome profiles. The user selects cells of interest in three interactive ways, and further gene-centric exploration is based on the selected subset of cells.

First, if annotation is present in the dataset, the user can select a cell type directly by selecting an annotation label from the data taxonomy.

Second, GeneSurfer is designed to seamlessly integrate with SpaceWalker^20^ by synchronizing the user-defined locations, where SpaceWalker automatically identifies transcriptomically similar cells at user-defined locations and GeneSurfer simultaneously provides further gene-centric exploration and GO term annotation within these cells. This has the advantage that cell selection is not restricted to prior discrete cluster boundaries, accommodating for a more gradual continuum of transcriptomic types.

Third, GeneSurfer can work with the dimensionality reduction plugin of ManiVault^24^. This workflow allows users to select cells from embeddings, such as HSNE^21^, t-SNE^11^ or UMAP^12^, computed on-the-fly from the data.

#### Gene filtering

Since the number of genes/features in scRNA-seq data can reach 30k^3^, we provide a filtering option before conducting gene clustering to remove uninformative genes. This step helps to selectively filter out genes that show specific spatial expression patterns or are highly expressed within the subset, ensuring a focus on the most informative genes for analysis.

The number of genes to keep for clustering is an interactively modifiable parameter. The user can dynamically observe the filtered genes displayed in the bar chart and a real-time overview of the clustered gene patterns while adjusting the desired number of genes. We provide three gene filtering modes: 1) differential expression between selected cells and the whole dataset, 2) Moran’s I, and 3) correlation of gene expression with spatial coordinate axes.

*Differential:* This option filters out genes that exhibit high expression among the selected cells, typically indicating cell type specific expression. Genes are sorted based on the difference in average expression levels between the selected cells and the whole dataset.

Moran’s I: Moran’s I^22^ measures spatial auto-correlation to identify genes with non-random expression patterns within the cell subset. Euclidean distances of the spatial coordinates are calculated to create a spatial weight matrix *w*. GeneSurfer then calculates Moran’s I for each gene using the weight matrix and gene expression within the cell subset as:

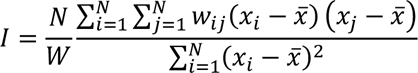

The calculated value is compared against the expected value under the null hypothesis:

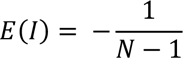

Z-score is calculated for each gene to indicate the statistical significance of the results. Genes are sorted by their Z-scores.

*Spatial:* This option is for uncovering significant expression correlations along specific spatial axes in the CCF, such as the dorsal-ventral or anterior -posterior axis. It calculates the Pearson correlation coefficient of each gene’s expression values with the spatial coordinates within the cell subset. Genes with high correlation scores are likely to exhibit strong expression patterns along the specified axes in the spatial domain.

#### Gene clustering

After filtering genes, hierarchical clustering^23^ is applied to group these filtered genes based on their local expression within the cell subset. The clustering algorithm uses a distance metric defined as 1 minus Pearson correlation score between the local expression of each gene pair. Subsequently, an average expression view of each cluster is displayed on the spatial map, providing a visual overview of gene expression patterns within each cluster. Meanwhile, filtered genes are color-coded in the bar chart according to their clustering labels.

Determining the optimal number of clusters has always been a challenge with clustering algorithms, especially in the context of data that is dynamically updated during exploration with our tool. Too many clusters can lead to overclustering, reducing the distinctiveness of gene expression patterns across clusters. Conversely, too few clusters might group genes with distinct patterns into one single cluster. To address this issue, we enable users to adjust the number of clusters on-the-fly, allowing for the determination of an appropriate cluster number through direct, iterative inspection for different data.

#### Gene set functional enrichment analysis

Upon selecting a gene cluster, gene set functional enrichment analysis is performed on-the-fly using the gProfiler^19^ or ToppGene^18^ API, and the results are dynamically updated with each cluster selection. The gene symbols of the selected gene cluster are sent to the chosen API and the enrichment analysis results are sent back to our system. The statistically significant GO terms, p-values and associated gene symbols are displayed in a table linked to the bar chart. Clicking on a GO term of interest highlights the associated genes in the bar chart, allowing for follow-up inspection of individual genes in 2D and 3D visualizations.

Users have the flexibility to switch between these APIs based on their preference. When using the gProfiler API, users can specify the species based on their dataset. For analysis of the limited ST gene set, in addition to sending the genes to be analyzed, all genes measured in the ST dataset are sent as background genes (500 genes for ABC Atlas). During exploration of whole-transcriptome scRNA data (> 30k genes for ABC Atlas), no background gene list is used.

#### Closed-loop interaction and parameter selection

A key contribution of GeneSurfer, distinguishing it from existing scripted transcriptomics data analysis methods, is its highly interactive interface designed specifically for exploratory analysis (see Supplemental Movie 1). The interface includes adjustable toolbars that allow users to tailor their data exploration. It offers direct, closed-loop visual feedback, showing users the effect of changes to hyperparameters.

Figure 6A shows a screenshot of the software interface. The interactive exploration loop starts with the selection of transcriptomically similar cells, as detailed in the *Selection of transcriptomically similar cells* section. Changes in the cell selection immediately trigger gene filtering and gene clustering based on user-defined settings, with options and adjustments detailed in the *Gene filtering* and *Gene clustering* section. Adjustments are made in real time using user interface controls, such as sliders and checkboxes, allowing users to see immediate updates in the bar chart of filtered genes and the clustering visualization of average gene expression. Clicking on a cluster triggers retrieval of the associated GO terms for genes in the selected cluster. The integration of cell selection, gene filtering and clustering, and visualization enables users to perform their analysis through a flexible, interactive interface.

**Figure 6.**
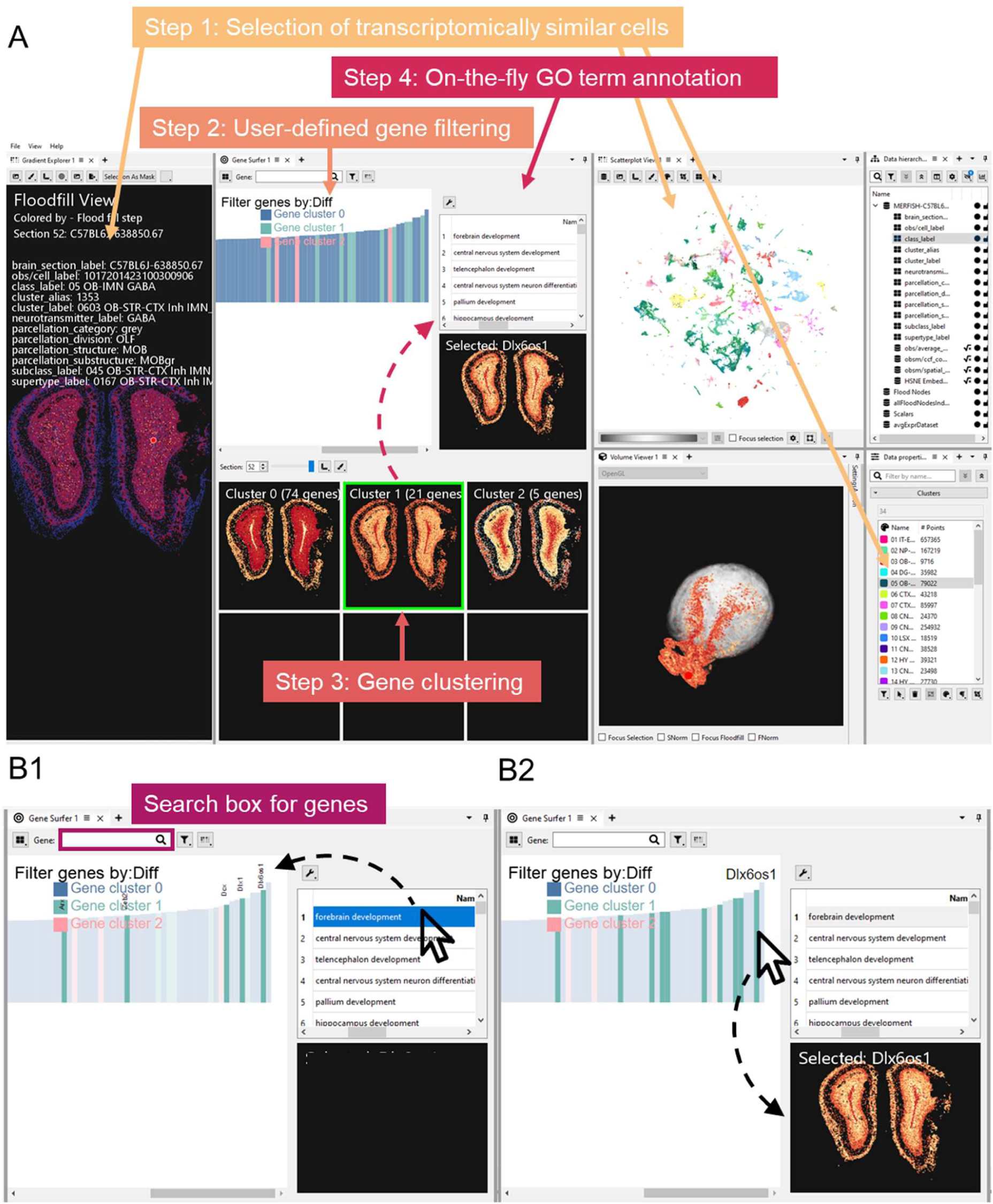
GeneSurfer interface and interaction loop. **A.** Screenshot of the interface of GeneSurfer with SpaceWalker. **B1.** When the user selects a GO term from the table, associated genes are highlighted in the bar chart. **B2.** Hovering over the bars shows the corresponding gene symbols; simultaneously, genes within the same cluster as the hovered gene are highlighted. Clicking on a gene displays its expression in both 2D and 3D visualizations.

One of our primary goals is to provide visualization of gene expression in the spatial domain, allowing users to inspect individual genes at each step of the exploration workflow. First, the user can visualize a specific gene in the dataset in 3D by entering the gene symbol in the search box (Figure 6B1). Second, selecting a GO term of interest from the annotation table highlights the associated genes in the bar chart (Figure 6B1). Third, hovering the mouse over the bars displays the gene symbol and highlights genes with the same cluster label. When the user clicks on a bar, the expression level of that gene is plotted in the spatial domain, enabling the user to select a gene of interest from the bar chart (Figure 6B2). This feature not only facilitates the exploration of genes associated with specific biological functions or pathways, but also allows for the inspection of individual gene expression in relation to selected GO terms.

In addition, point opacity for non-selected cells is adjustable from 0 to 1, providing flexibility in visualizing local expression. Setting the opacity to 0 isolates and displays only the selected cells, offering a focused view of the region of interest. Conversely, an opacity value of 1 renders all cells visible, providing a full overview of the spatial map. Intermediate opacity values allow for visual differentiation between the cells of interest and the surrounding cells, improving analytical clarity. Furthermore, the user can adjust the point size of the cells to adapt to the varying cell densities present in spatial datasets, enabling an optimal visualization across different scales of datasets.

## Supplemental Information

**Supplemental Movie 1. Demonstration of interactive exploration with GeneSurfer using ABC Atlas data.** The video showcases how users can select cells of interest, filter and cluster genes, retrieve associated GO terms and inspect gene expressions, for an interactive and dynamic analysis experience.

**Supplemental Table 1. Top 20 differentially expressed genes for each substructure compared to subclass 5310 Endo_NN_1.** This table accompanies Figure 5 and lists the top 20 genes that are differentially expressed in each substructure in the HSNE embedding compared to the endothelial subclass. Each column represents one substructure.

