## Supplemental Table 1 for "GeneSurfer Enables Transcriptome-wide Exploration and Functional Annotation of Gene Co-expression Modules in 3D Spatial Transcriptomics Data"

**Supplemental Table 1. Top 20 differentially expressed genes for each substructure compared to subclass 5310 Endo_NN_1.** This table accompanies Figure 5 and lists the top 20 genes that are differentially expressed in each substructure in the HSNE embedding compared to the endothelial subclass. Each column represents one substructure.

| 1 | 2 | 3 | 4 | 5 | 6 | 7 | 8 | 9 |
|---|---|---|---|---|---|---|---|---|
| *Slc17a7* | *Gpr88* | *Tcf7l2* | *Agt* | *Pvalb* | *Igf2* | *Gfap* | *Abcc9* | *Abcc9* |
| *Rasgrp1* | *Syndig1l* | *Shox2* | *Serpine2* | *Hopx* | *Dcn* | *Aqp4* | *Kcnj8* | *Kcnj8* |
| *Sv2b* | *Scn4b* | *Ramp3* | *Syt2* | *Calb1* | *Nr2f2* | *Agt* | *Rgs4* | *Egflam* |
| *Nfix* | *Penk* | *Tnnt1* | *Slc17a6* | *Slc38a1* | *Igfbp2* | *Lpar1* | *Egflam* | *Rgs4* |
| *Grm3* | *Bcl11b* | *Slc17a6* | *Unc13c* | *Ppp1r17* | *Slc7a11* | *Mog* | *Lhfp* | *Lhfp* |
| *Slc1a3* | *Drd2* | *Agt* | *Spon1* | *Shisa6* | *Gfap* | *Sox10* | *Ntn1* | *Ntn1* |
| *Adcy2* | *Crym* | *Spock3* | *Slc7a10* | *Kit* | *Egflam* | *Gja1* | *Slc17a7* | *Zic1* |
| *Nrn1* | *Spock3* | *Rasgrp1* | *Pvalb* | *Zic1* | *Pcp4l1* | *Hopx* | *Grm3* | *Nr2f2* |
| *Pou3f3* | *Rgs4* | *Spon1* | *Zfhx4* | *Unc13c* | *Lama1* | *Zeb2* | *Zic1* | *Igf2* |
| *Ptk2b* | *Caln1* | *Grm1* | *Gja1* | *Whrn* | *Fli1* | *Ccdc3* | *Nr2f2* | *Baiap3* |
| *Gda* | *Rasgrp1* | *Zic1* | *Aqp4* | *Ctxn3* | *Fxyd6* | *Slc38a1* | *Igf2* | *Gpc3* |
| *Pde1a* | *Drd1* | *Rgs4* | *Slc32a1* | *Slc1a3* | *Igfbp4* | *Marcksl1* | *Ror1* | *Agt* |
| *Cxcl14* | *Grm3* | *Lhfp* | *Gad2* | *Cnr1* | *Samd5* | *Bmpr1b* | *Car4* | *Ebf1* |
| *Rgs4* | *Gad2* | *Adra1b* | *Baiap3* | *Syndig1* | *Ntn1* | *Cdh20* | *Gpc3* | *Car4* |
| *Lrp4* | *Syt6* | *Nr2f1* | *Igsf1* | *Cbln1* | *Foxo1* | *Sox2* | *Tbx3* | *Ror1* |
| *St6galnac5* | *Slc32a1* | *Samd5* | *Cyp26b1* | *Syt2* | *Lhfp* | *Opalin* | *Ebf1* | *Grm3* |
| *Ccn3* | *Unc13c* | *Medag* | *Calb2* | *Grm1* | *Zfp521* | *Klk6* | *Rasgrp1* | *Unc13c* |
| *Ccnd2* | *Pcp4l1* | *Sema5a* | *Kcnk9* | *Ntn1* | *Maf* | *Chn2* | *Nr2f1* | *Tmie* |
| *Lamp5* | *Six3* | *Fam163a* | *Ramp1* | *Slc32a1* | *Vegfc* | *Prkcq* | *Samd5* | *Pde3a* |
| *C1ql3* | *Cnr1* | *Mctp2* | *Gsta4* | *Zfp521* | *Creb3l1* | *Dock5* | *Col27a1* | *Sema5a* |
